# Creation of Advance Construction of CRISPR-Cas9 via using Multi Round Gateway Cloning

**DOI:** 10.1101/2023.03.17.533097

**Authors:** M. Rizwan Jameel

## Abstract

CRISPR (Clustered Regularly Interspaced Short Palindromic Repeats)/Cas9 (CRISPR-associated system) is used to edit specific genomic sequences with precision and efficacy. CRISPR/Cas9 system mediated double strand binding site through error-prone non homologous end joining repair in targeted sequences, which the deletion or insertion can be achieved by knock out gene. Creation advance construct of CRISPR/Cas9 of soluble starch synthase II-1, 2 and 3 (*SSSII*) genes in single destination binary vector through multi round LR reaction of Gateway technology. Gateway cloning system developed the suitable for the site-specific DNA recombination of lambda-bacteriophage and *ccdB* the cytotoxic protein, which is poisonous to most E. coli strains. A genetic transformation vector designed with appropriate gRNAs, Cas9, and antibiotic resistance was used to create SSSII knockout mutants.

## INTRODUCTION

Clustered Regularly Interspaced Short Palindromic Repeats (CRISPR)/Cas9 endonuclease is a genomic editing tool. CRISPR/Cas9 is known for its simplicity, specificity, and versatility [1–5]. The functions of endonuclease Cas9 could be utilized to delete a specific site(s) of a gene using an engineered sequence single guide RNA (gRNA). The gRNA usually has 20 nucleotides against a specific target sequence in the genome [5, 6]. The SSS-II 1, 2, and 3 gene expression cassette was cloned into a plant transformation vector, pMDC99, employing the Multi round Gateway cloning techniques. The expression cassette pMDC99 was transferred into Indica rice through *Agrobacterium-mediated* genetic transformation. Mutations in soluble starch synthases (SSS) II [7–10] have altered the starch biosynthesis pathway in one way or the other in rice. Variation in SSS isoform profiles could affect the starch biosynthesis pathway and control phenotypes in the natural varieties of rice [11, 12]. Multi round Gateway technology is sitespecific recombinational cloning sysytem [14, 15]. This is site-specific recombination of bacteriophage λ through Gateway technology [15]. Bacteriophage λ assimilates keen on the chromosome of E. coli through site-specific recombination technique between the attachment sites (att) on the bacterial chromosome (attB) and the sites att of lying on the phage chromosome (attP), to produce left (attL) and right att sites (attR). This reaction of recombination is mediated via the integrase enzyme (Int) of bacteriophage λ and the E. coli integration host factor (IHF). The reaction of excision needs an extra host factor, excisionase (Xis), in adding to IHF and Int. The multi round Gateway cloning has developed changed att sites (such as attB1, attB2, attP1, and attP2) [14]. The attB1 sites specially recombine through the attP1 site, other than not attP2, to produce the attL1 and attR1 sites. Likewise, the attB2 site purposely recombines through the attP2 site to produce the attL2 and attR2 sites. In the initial step of the multi round Gateway cloning, the fragment of DNA for cloning is enlarged through primer sets of PCR having attB1 or attB2 sequences at the 5’ end points. Gateway technology cloning vectors have attP1 and attP2 sites for cloning. Consequently, the PCR product having the attB1 or attB2 sites at equally ends can be in particular inserted in between the attP1 and attP2 sites of the entry vector through sitespecific recombination system, to produce the entry clone of the Gateway technology [16]. The Gateway technology LR cloning reaction is mediated through the site-specific homologous DNA recombination properties of bacteriophage lambda [17]. This technique is more appropriate than other technique as it has not involved either DNA digestion or ligation, two processes this can hinder the cloning method [18]. The binary vector having a ccdB gene, which generates a toxic protein. It is lethal to mostly E.coli strains [19, 20]. In the Gateway cloning LR reaction, the *ccdB* gene in the binary vector is substituted through the targeted gene from an entry vector with site-specific DNA homologous recombination. The multi round LR reaction mixture having both recombinant and non-recombinant DNA plasmids are consequently transmitted keen on E. coli bacteria, such as pMDC99, which is sensitive to the toxic effect of *ccdB.* Only recombinant pMDC99 plasmids that have lost the ccdB gene are capable to stay alive in the transformed EHA105 cells. Lately, collections of Gateway technology compatible binary destination vectors, that can be used for *Agrobacterium* mediated transformation, have been created and are obtainable to the plant research community [21–24]. Targeted sequences can be simply cloned into these binary destination vectors throughout Gateway technology multi round LR cloning [25]. The main Gateway cloning technique was additional enhanced for cloning multiple DNA fragments into a single vector [26, 27]. Recently, up to four targeted gene sequences can be cloned into a single vector destination. The 4-fragment MultiSite Gateway cloning technology is suitable for the construction of a donor DNA plasmid for CRISPR/Cas9 and NHEJ-mediated gene knocks out [16].

## MATERIAL AND METHODS

The bacterial host strains, chemicals, cloning vectors, enzymes and planting materials, etc. utilized for the gene cloning, protein expression and purification, plant transformation, transgenic screening, and Starch assessment and their sources.

### Primer design

The primers were designed using Mac-Vector (Acceleris, GmbH), Snap Gene (form 1.1.3), or using electronic programming, for example, Primer blast (http://www.ncbi.nlm.nih.gov/instruments/groundworkimpact/list) and Primer 3 plus (https://primer3plus.com/) with default values. Oligo-analyzer programming (http://www.idtdna.com) was used to check for the probability of self or hetero-dimer development in the planned set of Primers. These Primers were confirmed for their uniqueness using BLASTN at the NCBI database. The primers were further combined using IDT, Germany, or Sigma-Aldrich, India [6].

### gRNA Synthesis of advance Construct of SSSIIs genes

The target site to be knocked out was picked with the end goal that it lays in the exonic region just as it is the exon just which partakes in gene expression. Thus, to pick the target sites the cDNA sequence of the gene of intrigue was utilized. Gene editing in the intronic region would have no impact on gene working. In CRISPR innovation, just the gRNA shifts while the remainder of the cassettes, for example, scaffold, promoter, and vector spine continues as before. The gRNA could be changed by necessity only by digesting the scaffold vector with an appropriate restriction enzyme to clone sgRNA in it. To predict imminent gRNAs for both the gene, the cDNA succession of all three *OsSSSII-1, OsSSSII-2, and OsSSSII-3* genes was taken in online CRISPR-direct programming. Checking sense and antisense strands of both the cDNAs, the CRISPR direct programming anticipated a few imminent target sites which were promptly adjoining PAM sequences. Consequently, the target sequence to be mutated resembled 5’-N (20) - NGG-3’ or 5’-CCN-N (20)- 3’. Our target sequence which is 20 ntds, was available only nearby the PAM sequence that is 5’NGG3’. Both the forward and turn around single-stranded oligos of 20 ntds were incorporated from IDT (USA). The complementary forward and turn around oligos of all gRNAs were toughened to one another in thermo-cycler to frame a double-stranded gRNA having four ntds. Shades on either side, which is correlative to the shades of processed EV1 and EV2. Two separate articulation vectors were intended for targeting *OsSSSII-1, OsSSSII-2, and OsSSSII-3* genes [12].

### To Design advance Construct

The cloning technique included the utilization of three gateway cloning viable vectors i.e., entry vector1 (EV1), entry vector2 (EV2), and Destination vector (DV), likewise called an expression vector. Right off the bat, the gRNA1 and gRNA2 were cloned into the Bsa1 site of EV1 and EV2 entry vectors, separately. At that point, *SSSII-1* gRNA2, *SSSII-2* gRNA1 and *SSSII-3* gRNA2 from these entry vectors were cloned into Destination vector (previously containing Cas9 gene cassette) individually through gateway cloning utilizing LR clonase. The last Construct got was changed into Agrobacterium EHA105 cells through electroporation technique [12].

### Multi Round Gateway cloning

The Gateway technology is a dynamic and all-inclusive cloning technology dependent on the guideline of homologous site-explicit recombination which helps in the mix of lambda (λ) phage into the E. coli chromosome [28]. The recombination of λ phage requires site-explicit attachment (att) sites: attP on the λ phage chromosome and attB on the E. coli chromosome. The sites attachment are all around described and discovered to be vital for the binding of recombination proteins [29]. The λ recombination is bio-catalyzed by various compounds that target and tie to the attachment (att) sites, join the target sites, limit them, lastly covalently attach the DNA. Within the sight of λ integrase and excisionase, homologous recombination happens between viable att sites [30]. Utilizing the λ recombination system-based Gateway cloning innovation it is possible to move heterologous DNA sequences (flanked by altered att locales) between vectors without the need of restriction catalysts [31]. The Multi Round Gateway cloning technology [32] considers the in vitro gene pyramiding/stacking of various genes into a solitary plant change vector through a series of recombination responses.

### The steps engaged with LR Multi-Round Gateway cloning technologies are as per the following

- The initial step includes endonuclease based cloning of the cassettes gene expression into plasmids having flanking “att L” recombination sites to create “Section Vectors” (EV-1 and EV-2)
- The second step includes the exchange of the cassette gene in the Gateway Entry Vector into a Gateway Destination vector plasmid that contains “att R” recombination sites (pMDC99) utilizing an exclusive enzyme blend, “LR Clonase™” (Invitrogen, USA).

With the end goal of Multi-Round Gateway cloning two diverse entry vectors, EV-1 and EV-2 were utilized. These two entry vectors contrast in the recombination sites, EV1 (pL12R-34H-Ap) encloses the recombination connection sites attL1-attL2 and attR3-attR4 though, EV2 (pL34R12H-CmR-ccdB) contains the recombination sites attL3-attL4 and attR1-R2. These entry vectors (EV1 and EV2) can be utilized related to the gateway viable destination vectors (pMDC99-plant change vector) conveying the attR1-attR2 recombination destinations for the development of multi-quality plant change builds. The pMDC99 vector contains the hygromycin obstruction gene hygromycin phosphotransferase *(hpt)* as a plant determination marker [33].

### LR cloning of Mutli round Gateway Cloning

The LR recombination of the first round consisted of recombination between EV-1 recombinant vector (containing the MzUbip: Cas9: NosT) and therefore the pMDC99 destination vector. The recombinant first LR inclusion (Cas9:pMDC99) was isolated from the positive colonies to be used for the LR cloning of the second round. The LR second round reaction implanted the recombination between *SSSII-1* gRNA2 recombinant vector (having gRNA2 expression cassette: EV2:U6promoter:gRNA2: scaffold) and therefore the destination vector plasmid (pMDC99:Cas9) obtained once initial LR reaction. The LR third round recombination consisted of the recombination reaction between *SSSII-2* EV1 recombinant vector (having the gRNA1 expression cassette: U3promoter: gRNA1: scaffold) and the pMDC99:Cas9:gRNA2 vector. The LR fourth round recombination consisted of the recombination reaction between *SSSII-3* EV2 recombinant vector (having the gRNA2 expression cassette: U6promoter: gRNA2: scaffold) and therefore the pMDC99:Cas9:gRNA2:gRNA1 vector. The LR fourth round recombination consisted of the recombination reaction between *SSSII-3* EV2 recombinant vector (having the gRNA2 expression cassette: U6promoter: gRNA2: scaffold) and the pMDC99:Cas9:gRNA2:gRNA1 vector. The LR final round recombination consisted of the recombination reaction between EV1 vector (having the gRNA1 expression cassette) and the pMDC99:Cas9:gRNA2:gRNA1:gRNA2 vector to remove *ccdB* gene from advance vector of pMDC99:Cas9: *SSSII-1* gRNA2: SSSII-2 gRNA1: SSSII-3 gRNA2.The EV-2 vector of *OsSSSII-1,* EV-1 vector of *OsSSSII-2 and* EV-2 vector of *OsSSSII-3,* respective genes containing the gRNA2, gRNA1 and gRNA2 expression cassette was combined with pMDC99:Cas9:gRNA2/gRNA1/gRNA2 vector during a 2:1 quantitative relation (nanomoles) within the presence of LR clonase catalyst mix and incubated at 16 °C for 19 hours. This LR reaction was terminated by adding one μl enzyme K. associate degree aliquot of 2μl of the primary round LR reaction product was remodeled into E. coli top10/ E. coli DB3.1 (EV2/gRNA2 is related to E. coli DB3.1) cells exploitation heat shock and designated on lb agar plates supplemented with 50 μg/ml kanamycin. The resistant colonies obtained were screened for the presence of a gRNAs expression cassette by colony PCR exploitation sequence-specific primers. The LR final round cloning was transferred into *Agrobacterium* (EHA105) via electroporation and chosen on YEM agar plate supplemented with 50 μg/ml kanamycin, 50 μg/ml chloramphenicol and 50 μg/ml rifampicin. The ultimate expression vector construct (pMDC99:Cas9: *OsSSSII-1-gRNA1*:gRNA2, pMDC99: Cas9: *OsSSSII-2-* gRNA1: gRNA2 and pMDC99: Cas9: *OsSSSII*-3-gRNA1: gRNA2) was checked for the presence of *hpt* gene, Cas9 gene, gRNA1, and gRNA2 discrimination specific sets of screening primers. Delineated illustration of ultimate expression vector having Cas9, gRNA expression cassette. The sequence of screening primers of hygromycin, Cas9 genes, gRNAs of all three *OsSSSII-1, OsSSSII-2 and OsSSSII-3* Expression.

**Table 1:**
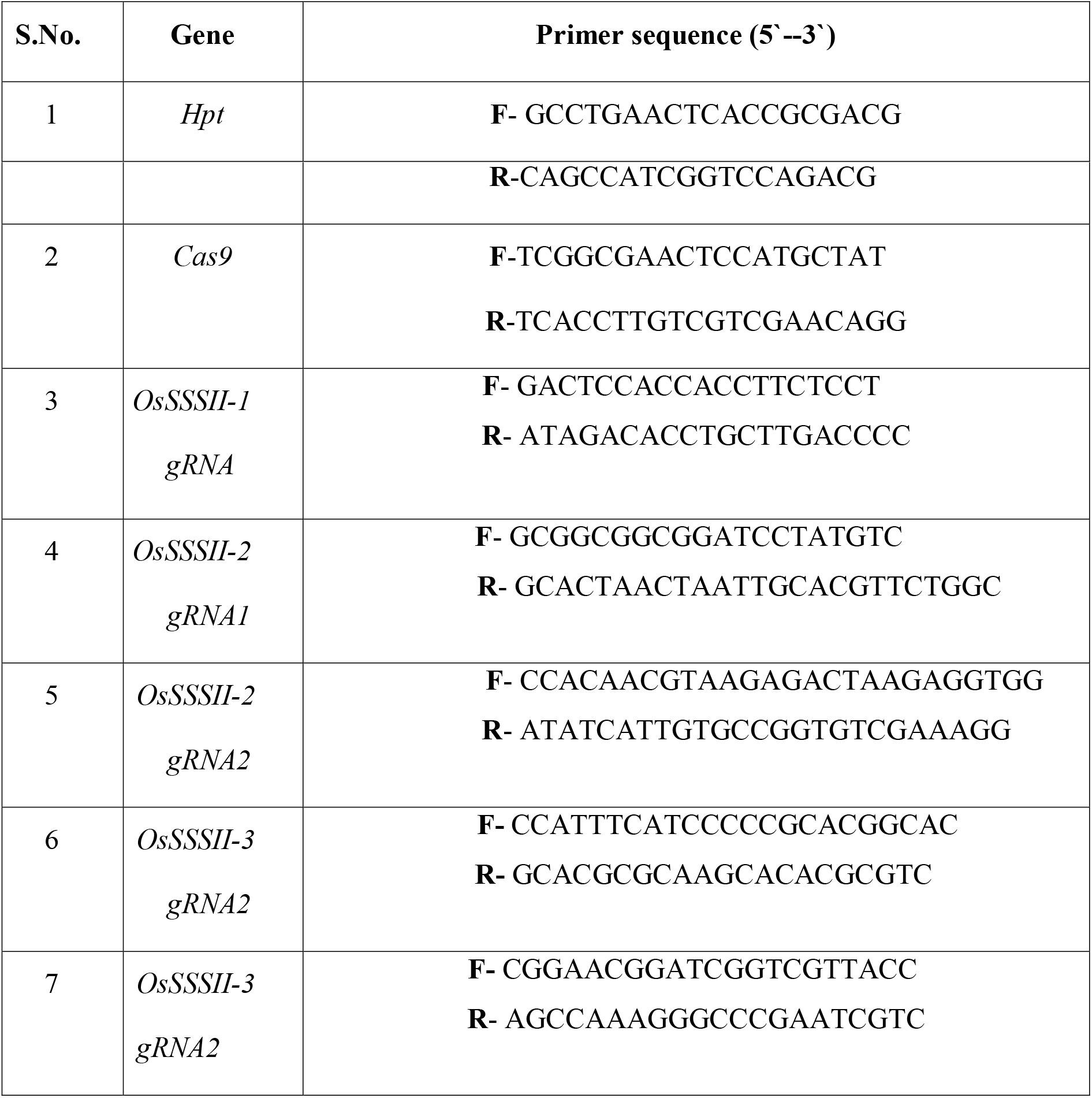
Primers of sgRNAs sequences for targeted genes [12].

## RESULT

### 1. The Digestion of EV1 and EV2 with Bsa1 Endonuclease

Both vectors EV1 and EV2 were digested with Bsa1 endonuclease to facilitate clone gRNA cloning. The digested fragments were run on 1% agarose gel. The EV1 vector provided 1 band of figure 2 and figure 3 sizes of ~2.5 kb and around 750 bp. The 3 kb vector was slash from the agarose gel and purified through quigen kit and after that utilized to clone gRNA2 in it by means of the Bsa1 site. In the same way, the EV2 digested through Bsa1 gave 2 band of figure 2 size of ~4 kb and 350 bp outcomes. The Bsa1 digestion of Entry vectors through usual size outcome is shown in gel figure 2 and figure 3.

**Figure 1:**
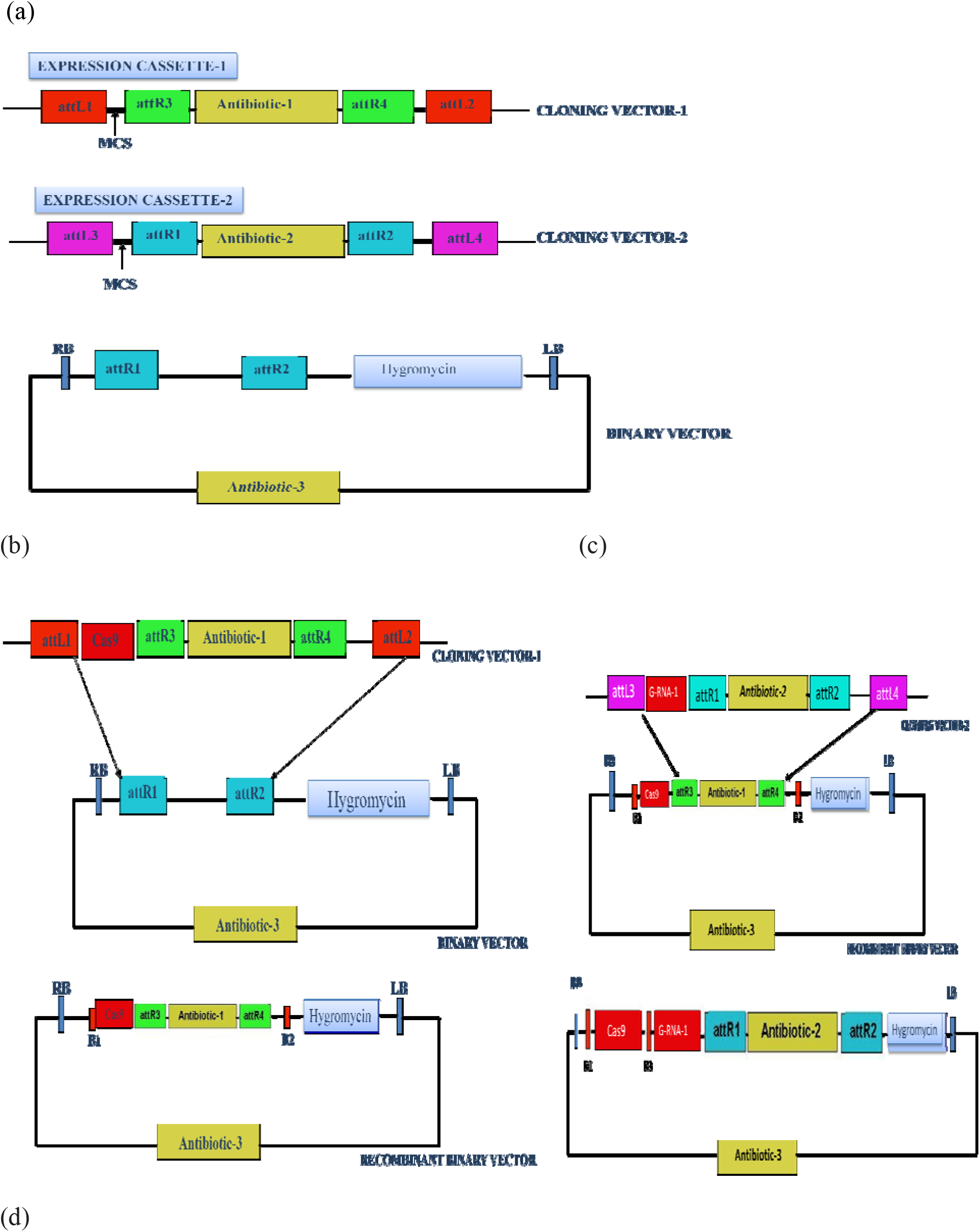
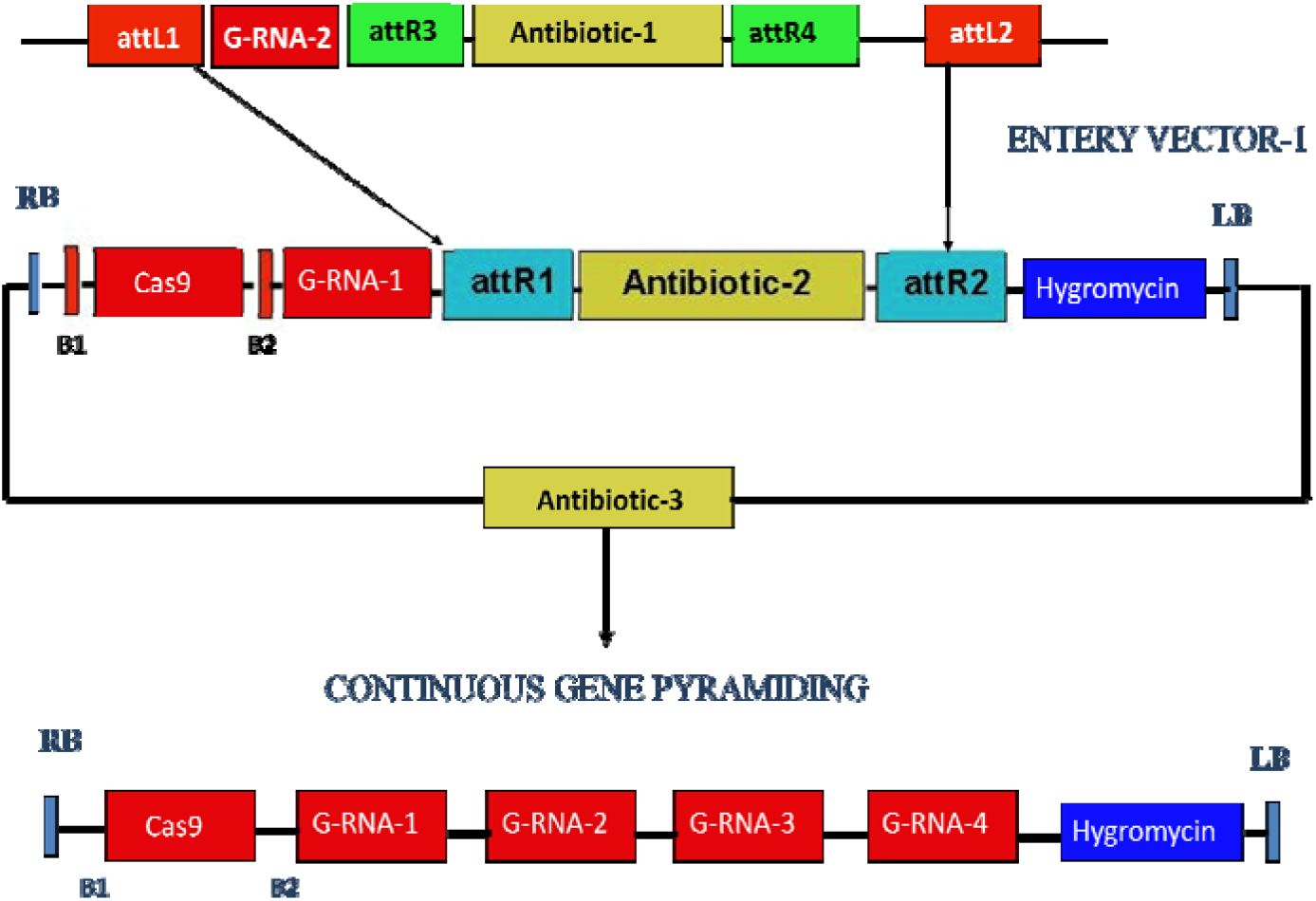
Sequential Multi gateway Cloning Vectors for Indefinite Stacking of genes (a) First round of Stacking, (b), second round of Stacking (c) and third round of Stacking **(d)**

**Figure 2:**
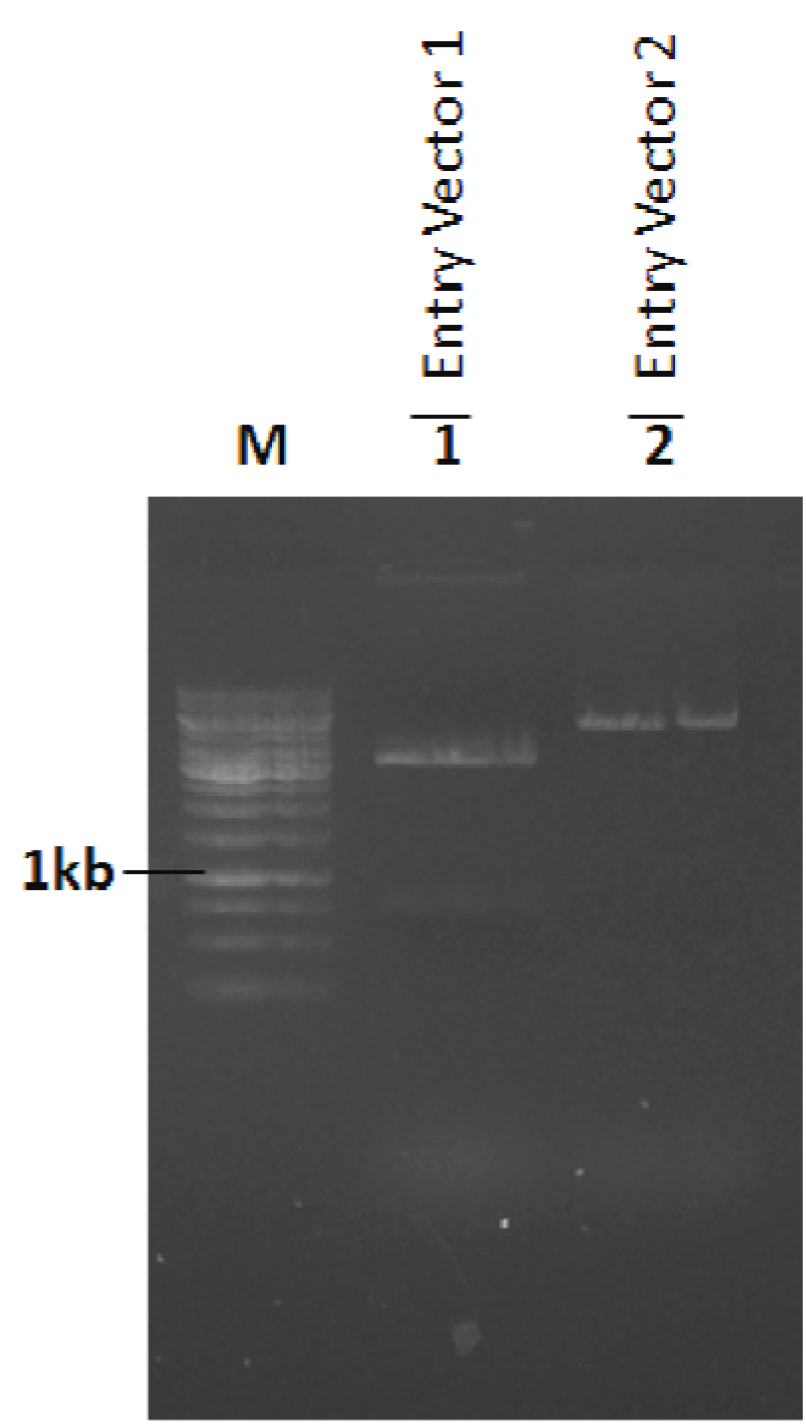
Plasmid digestion of Entry Vectors with BSA I. The band M is 1 kb ladder, band 1 is Entry Vector 1 and band 2 is Entry Vector 2.

**Figure 3:**
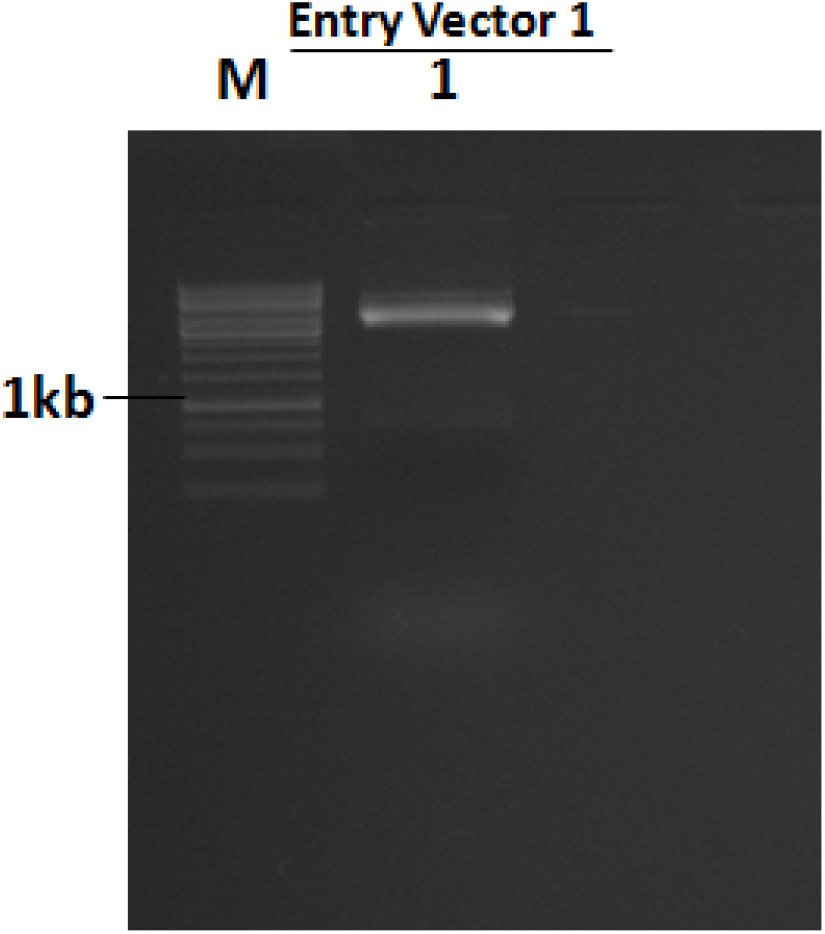
Plasmid digestion of Entry Vector with BSA I. The band M is 1 kb ladder and band 1 is Entry Vector 1.

### 2. Confirmation of gRNAs in recombinant vectors EV1 and EV2 with using PCR of advance Construct of SSSIIs genes

The recombinant vector EV1 containing promoter U3 and gRNA1 of one *SSSII-2* gene cloned in it was changed into TOP10 cells and plated on LB media enhanced with gentamycin for the determination of transformed. The putative changed states having EV2; *SSSII-1*-gRNA2 and EV2; *SSSII-3*-gRNA2 were screened through colonies PCR so as to check the presence of gRNA2 utilizing forward primers of U6 promoter and reverse primer of gRNA2. The PCR items from the gRNA1 transformants gave a normal size band of ~ 670 bp. State PCR of both the EV1 gave a normal size band; which affirms the right get together of gRNA1 cassettes in the EV1. In the same way, the presumed transformed DB3.1 cells containing recombinant vector EV2 that is EV2; *SSSII-1*-gRNA2, EV1; *SSSII-2*-gRNA1 and EV2; *SSSII-3-*gRNA2 was confirmed throughout colony PCR where forward primer of promoter U3 and reverse primer of gRNA1 was utilized. The ensuing PCR amplicon for all of two EV2; *SSSII-1*-gRNA2 and EV2; *SSSII-3-* gRNA2 was of ~750 bp. Hence, the gRNA1 was effectively cloned in EV1. Three ones the plasmids recombinant was cut off from transformed cells for extra use in LR multi-round gateway cloning. In the same way, Entry vector 1 and Cas9 gene expression cassette was ensured with colony PCR with help specific set of primers of maize ubiquitin promoter, Cas9 gene, and NosT. The PCR amplification are run 1 % agarose gel and developed bands in the gel are in the gel figures 4 to 17

**Figure 4:**
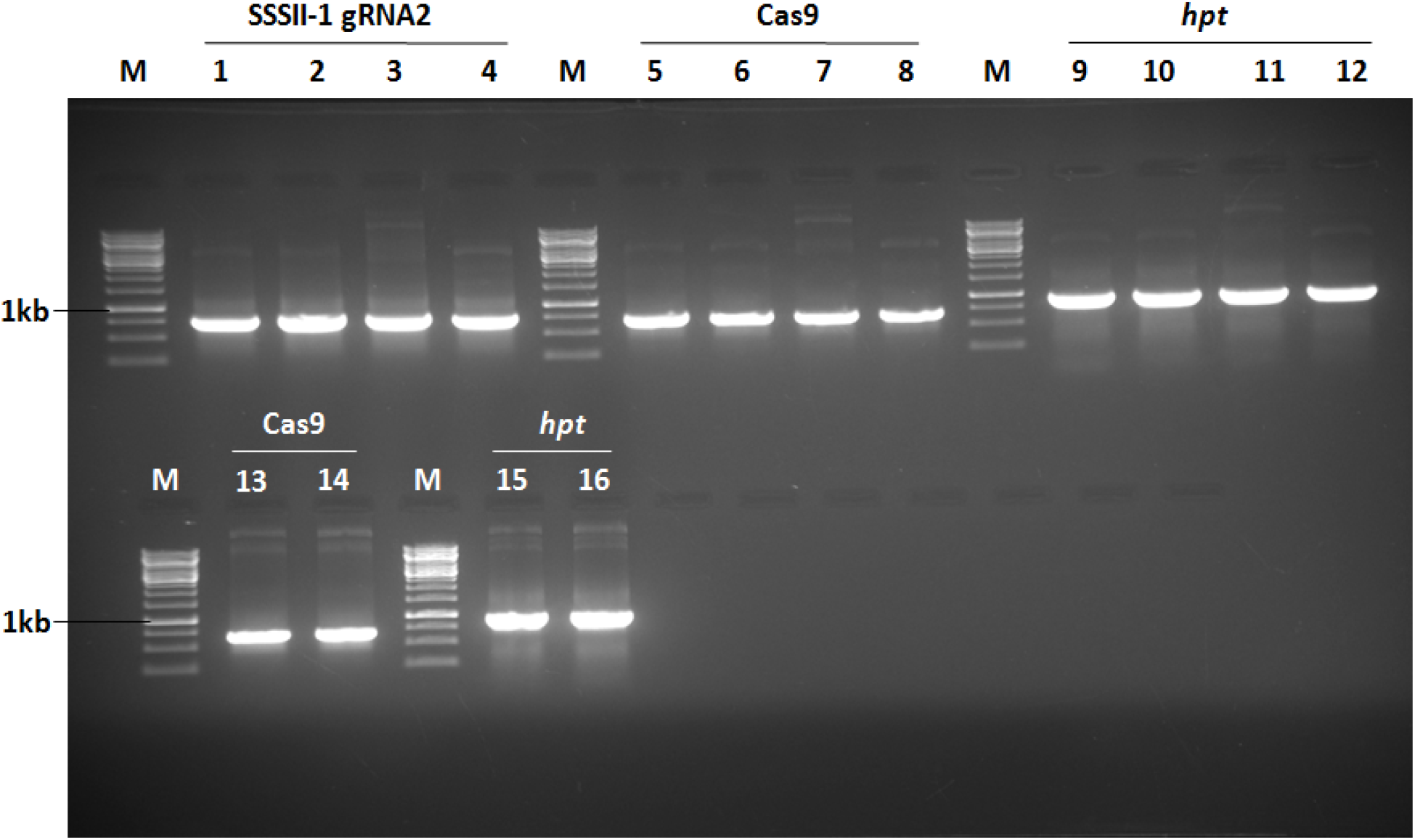
The formation of advance construction of all three isoforms of soluble starch synthases: SSSII-1, 2, and 3 in the single binary vector. The Band M, 1 kb DNA ladder (Invitrogen, MA, USA). The confirmation with PCR of bands 1 to 4 for SSSII-1 target 2 primer, bands 5 to 8 of Cas9 primers, bands 9 to 12 of *hpt gene* primers, also bands 13 and 14 of Cas9, and bands 15 and 16 of *hpt* gene primers.

**Figure 5:**
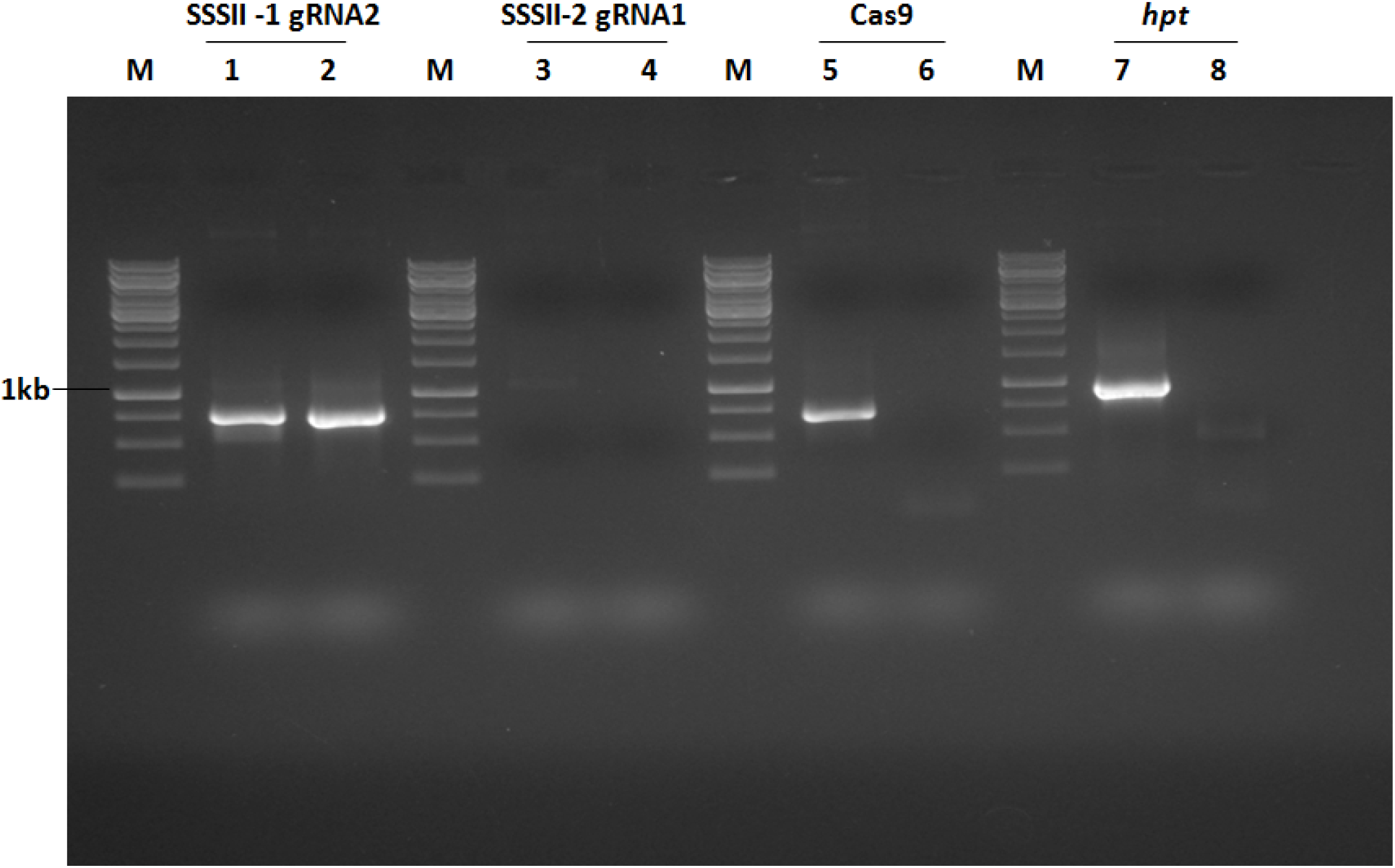
The formation of advance construction of all three isoforms of soluble starch synthases: SSSII-1, 2, and 3 in single destination vector. The Band M, 1 kb DNA ladder (Invitrogen, MA, USA). The confirmation with PCR of bands 1 to 2 for SSSII-1 target 2 primer, bands 3 to 4 for SSSII-2 target 1 primer bands 5 to 6 of Cas9 primers and bands 7 to 8 of *hpt gene* primers.

**Figure 6:**
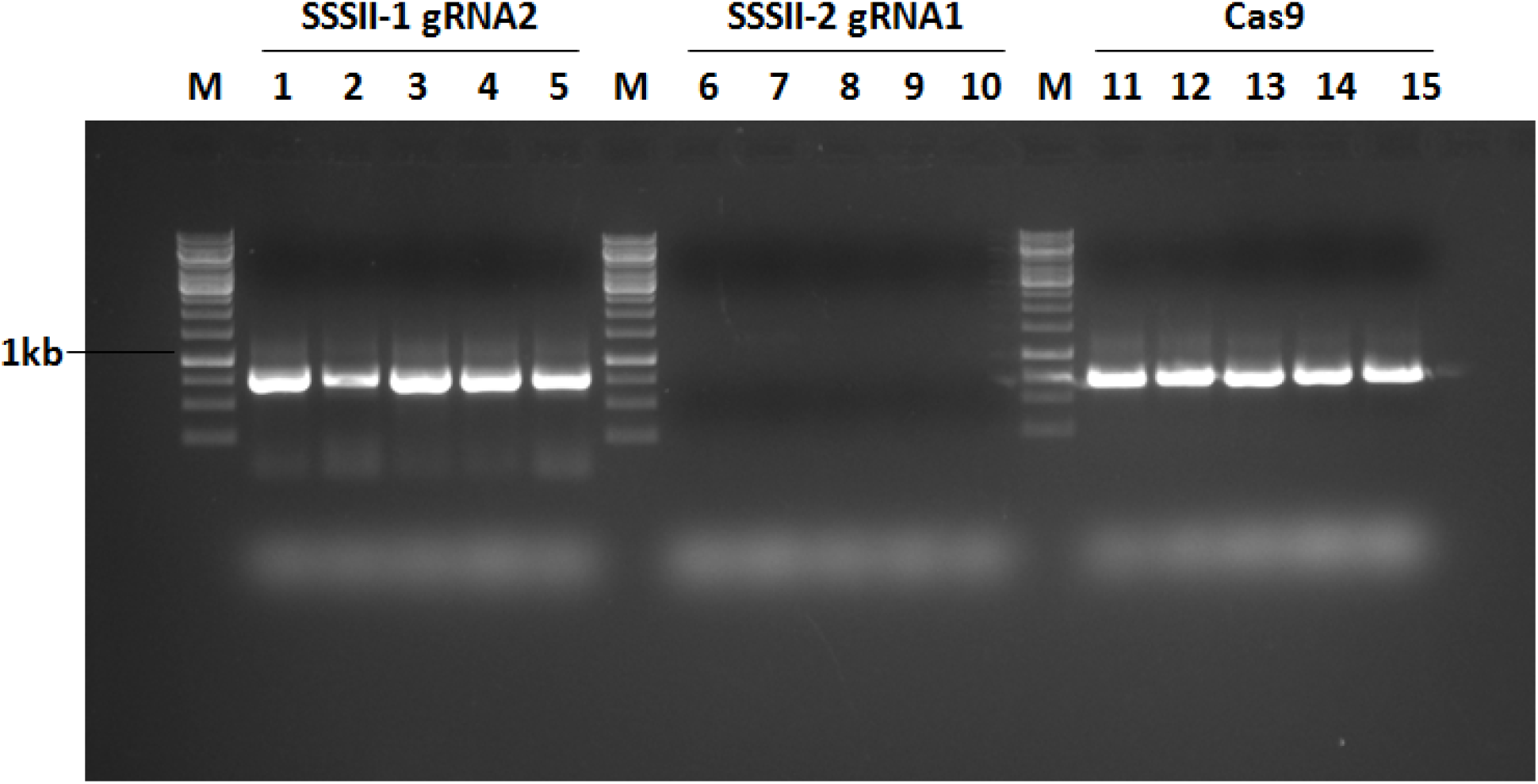
The formation of advance construction of all three isoforms of soluble starch synthases: SSSII-1, 2, and 3 in single destination vector. The Band M is 1 kb DNA ladder. The confirmation with PCR of bands 1 to 5 for SSSII-1 target 2 primer, bands 6 to 10 for SSSII-2 target 1 primers and bands 11 to 15 of Cas9 primers.

**Figure 7:**
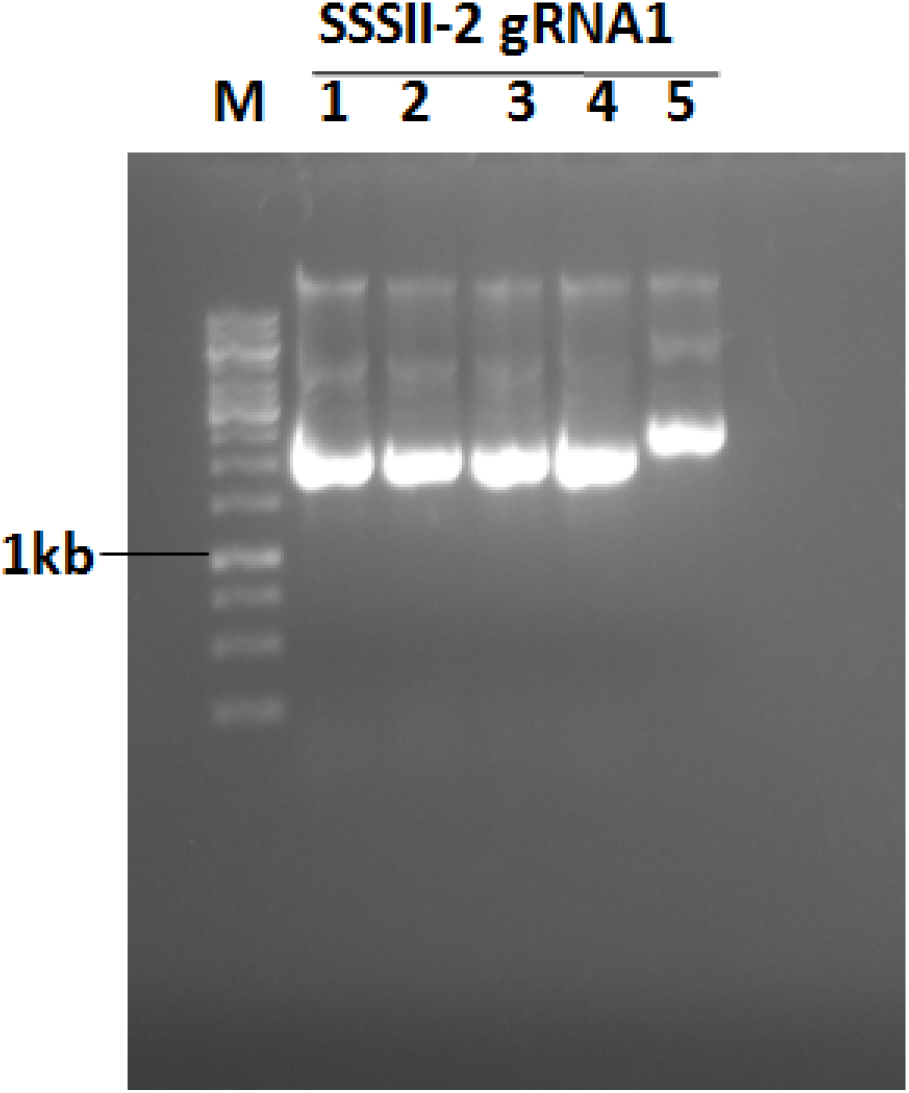
The formation of advance construction of all three isoforms of soluble starch synthases: *SSSII-1, 2, and 3* in single destination vector. The Band M is 1 kb DNA ladder. The confirmation with PCR of bands 1 to 5 for *SSSII-1* target 2 primers.

**Figure 8:**
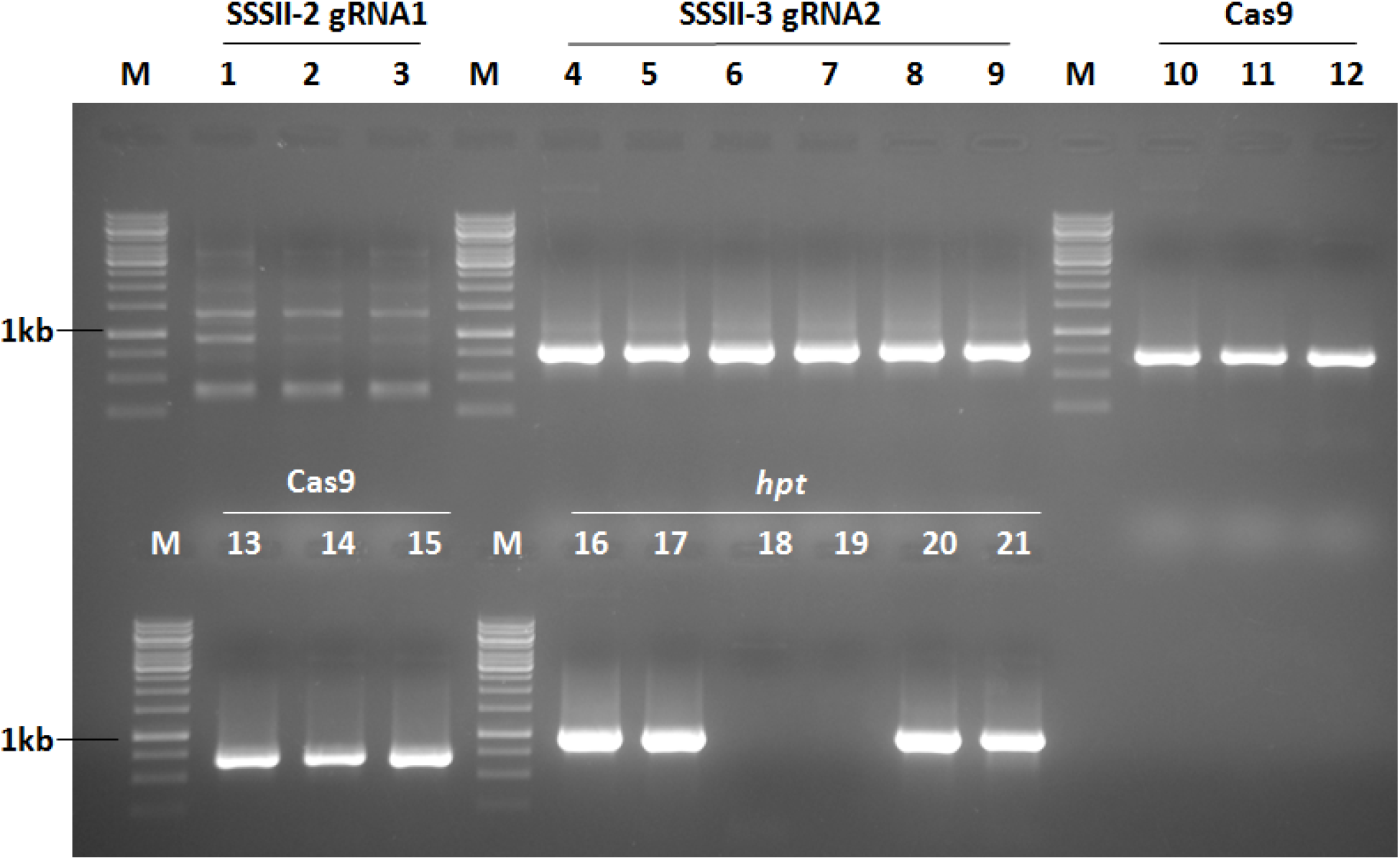
The formation of advance construction of all three isoforms of soluble starch synthases: SSSII-1, 2, and 3 in single destination vector. The Band M is 1 kb DNA ladder. The confirmation with PCR of bands 1 to 3 for SSSII-2 target 1 primer, bands 4 to 9 for SSSII-3 target 2 primers, bands 10 to 15 of Cas9 primers and bands 16 to 21 of *hpt gene* primers.

**Figure 9:**
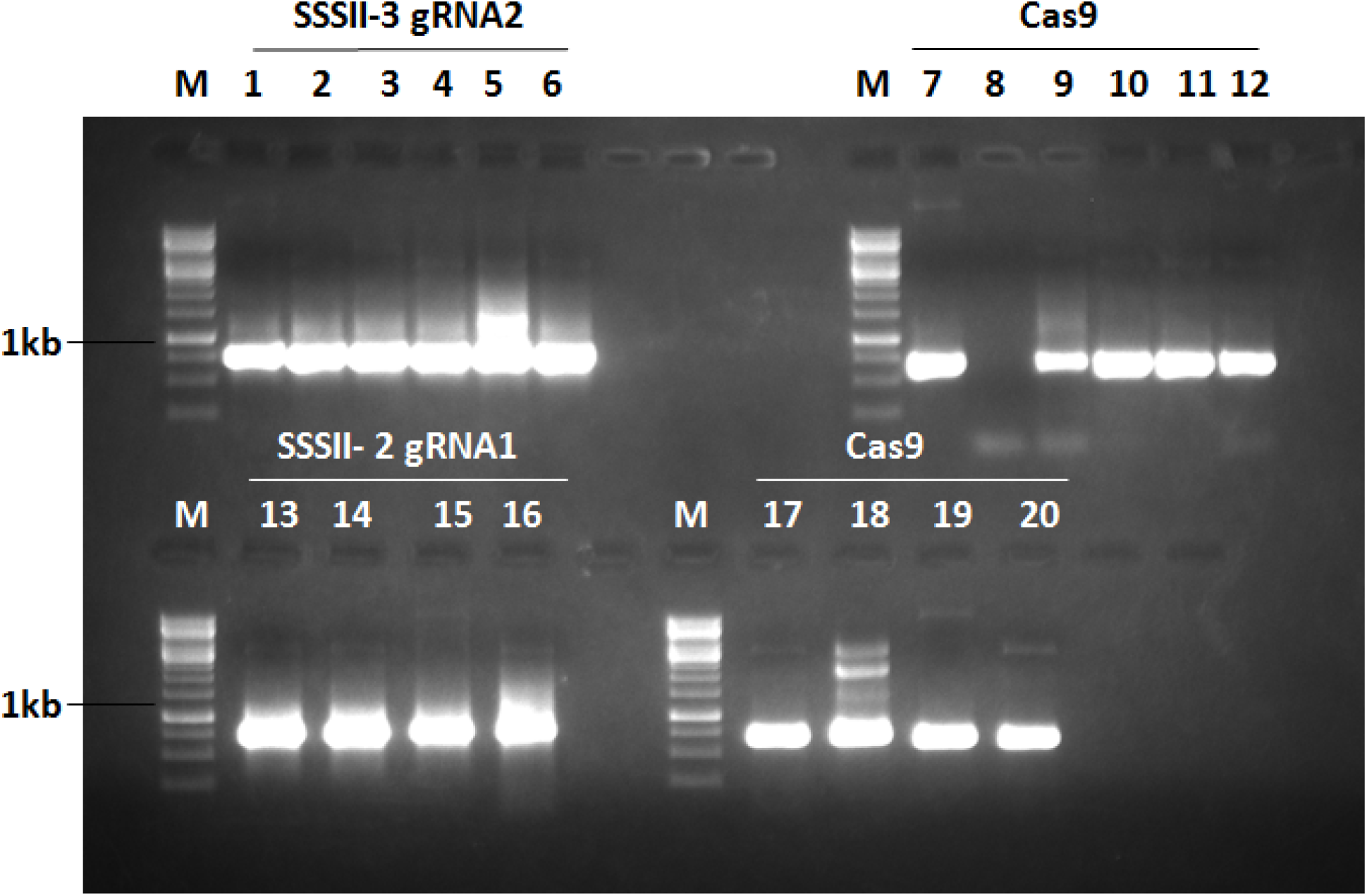
The formation of advance construction of all three isoforms of soluble starch synthases: SSSII-1, 2, and 3 in single destination vector. The Band M is 1 kb DNA ladder. The confirmation with clonies PCR of bands 1 to 6 for SSSII-3 target 2 primer, bands 7 to 12 of Cas9 primers, bands 13 to 16 for SSSII-2 target 1 primers and bands 17 to 20 of *hpt* primers.

**Figure 10:**
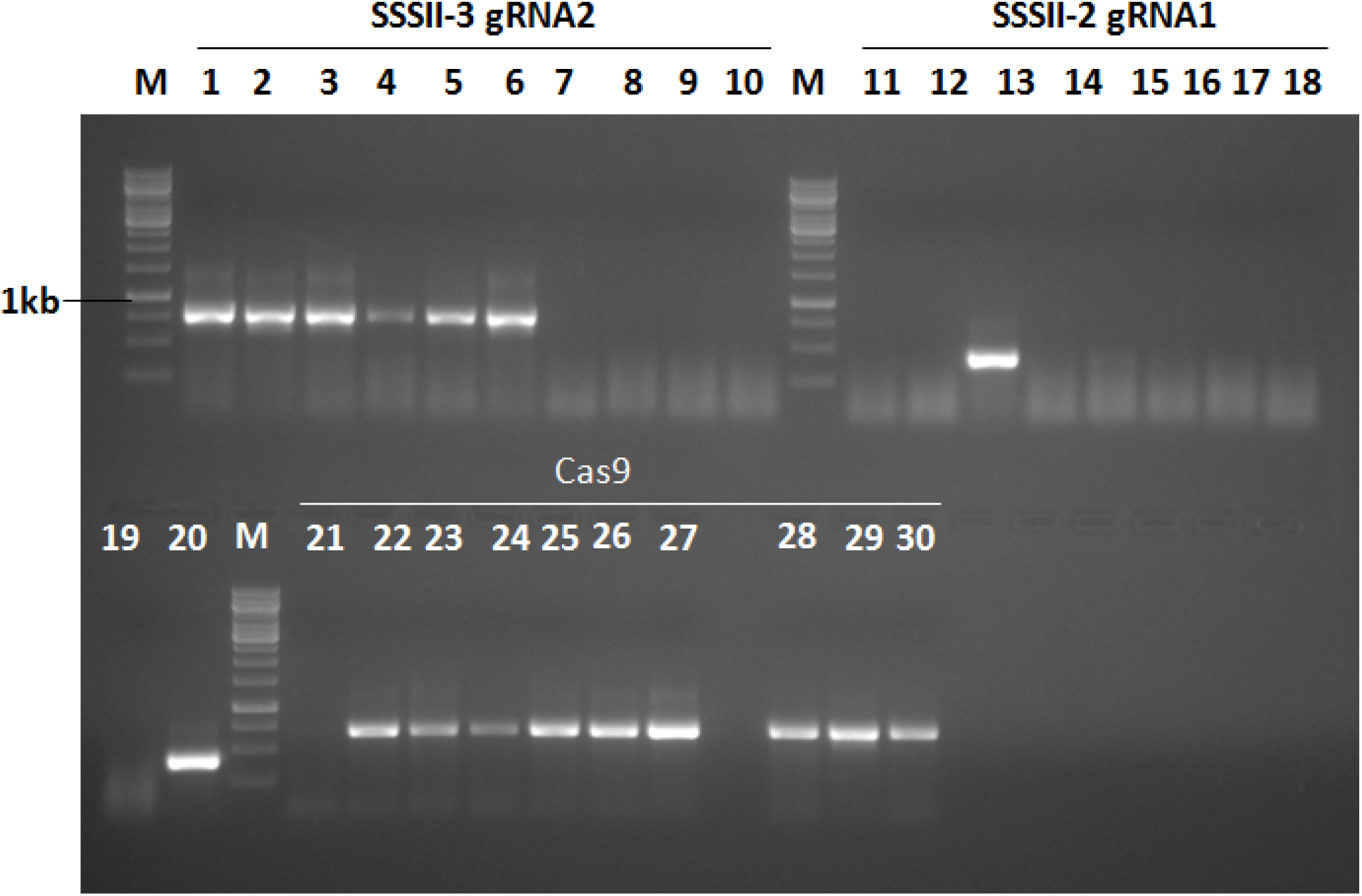
The formation of advance construction of all three isoforms of soluble starch synthases: *SSSII-1, 2, and 3* in single destination vector. The Band M is 1 kb DNA ladder. The confirmation with PCR of bands 1 to 10 for SSSII-3 target 2 primers, bands 11 to 20 for SSSII-2 target 1 primers and bands 21 to 30 for Cas9 primers.

**Figure 11:**
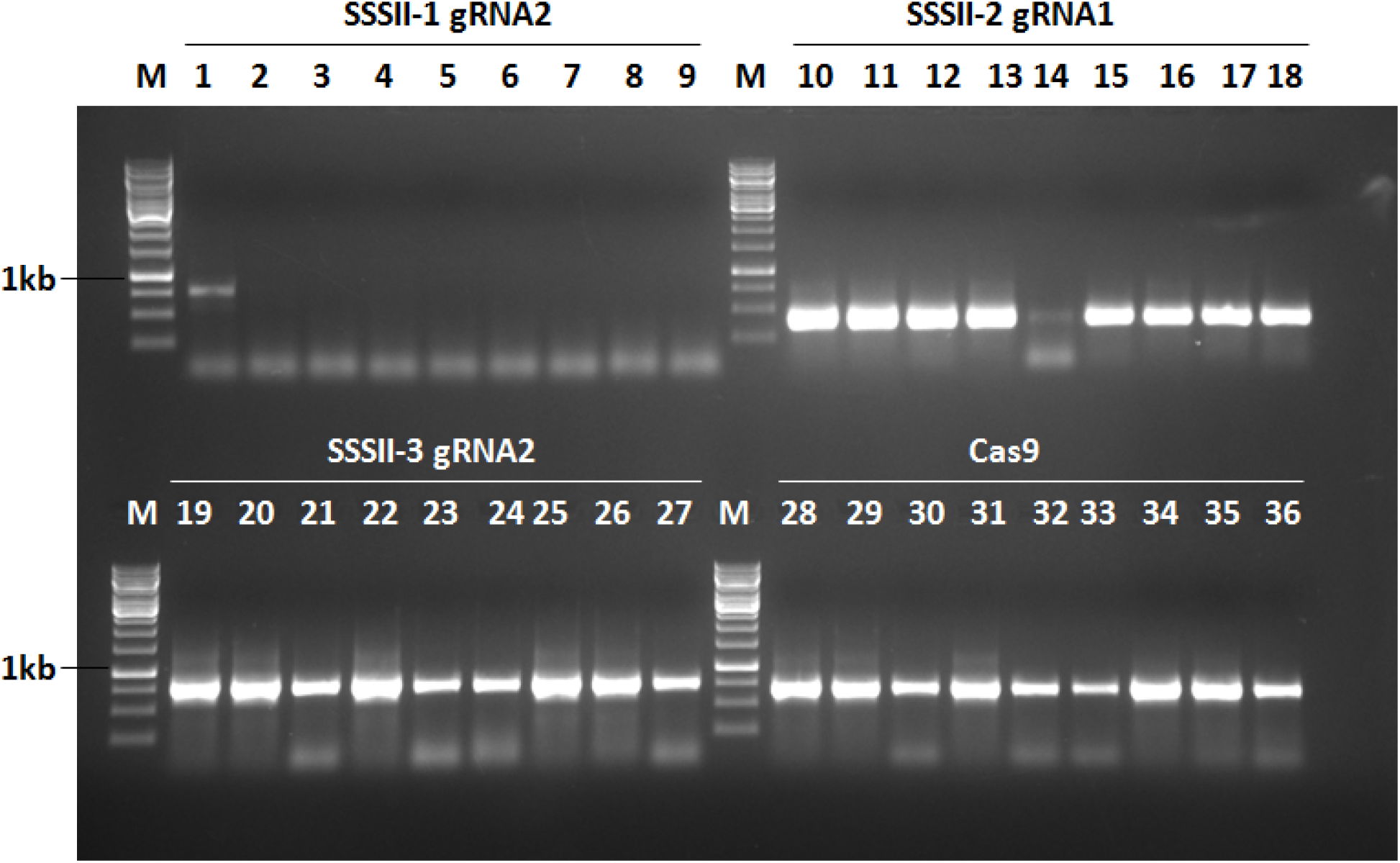
The formation of advance construction of all three isoforms of soluble starch synthases: SSSII-1, 2, and 3 in single destination vector. The Band M is 1 kb DNA ladder. The confirmation with PCR of bands 1 to 9 for SSSII-1 target 2 primers, bands 10 to 18 for SSSII-2 target 1 primers, bands 19 to 27 for SSSII-3 target 2 primers and bands 28 to 36 for Cas9 primers.

**Figure 12:**
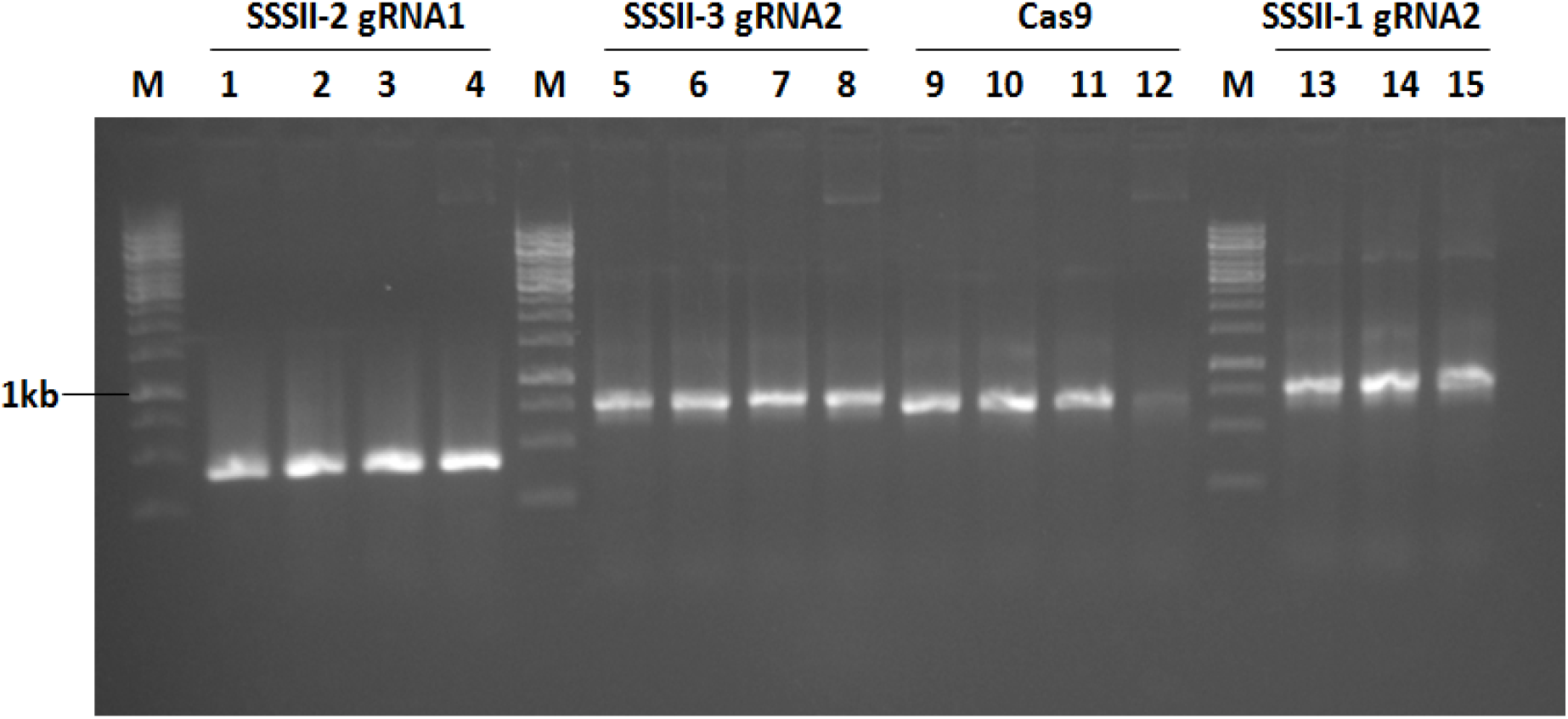
The formation of advance construction of all three isoforms of soluble starch synthases: SSSII-1, 2, and 3 in single destination vector. The Band M is 1 kb DNA ladder. The confirmation with PCR of bands 1 to 4 for SSSII-2 target 1 primers, bands 5 to 8 for SSSII-3 target 2 primers, bands 9 to 12 for Cas9 primers and bands 5 to 8 for SSSII-1 target 2 primers

**Figure 13:**
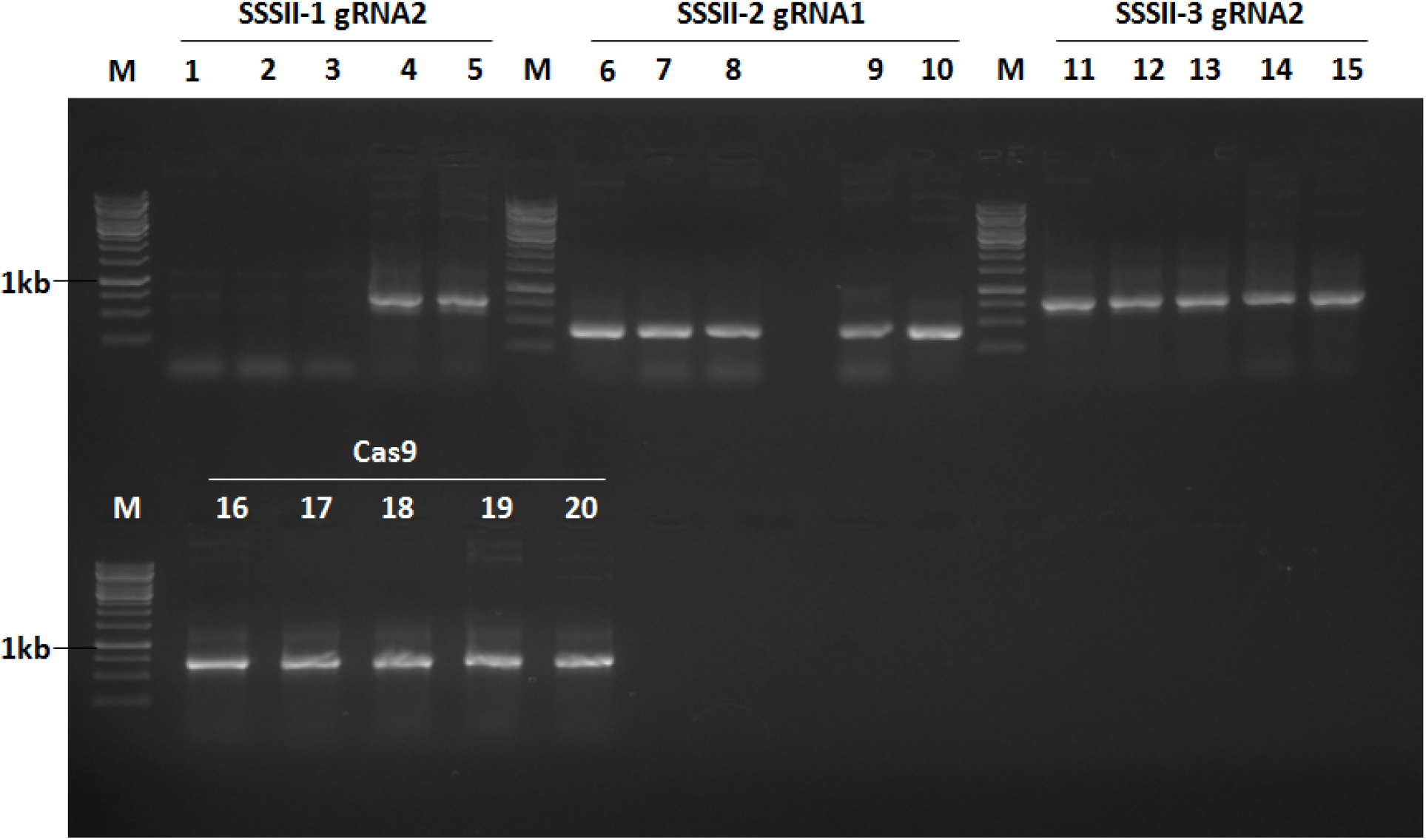
The formation of advance construction of all three isoforms of soluble starch synthases: SSSII-1, 2, and 3 in single destination vector. The Band M is 1 kb DNA ladder. The confirmation with PCR of bands 1 to 4 for SSSII-1 target 2 primers, bands 6 to 10 for SSSII-2 target 1 primers, bands 11 to 15 for SSSII-3 target 2 primers and bands 16 to 20 for Cas9 primers.

**Figure 14:**
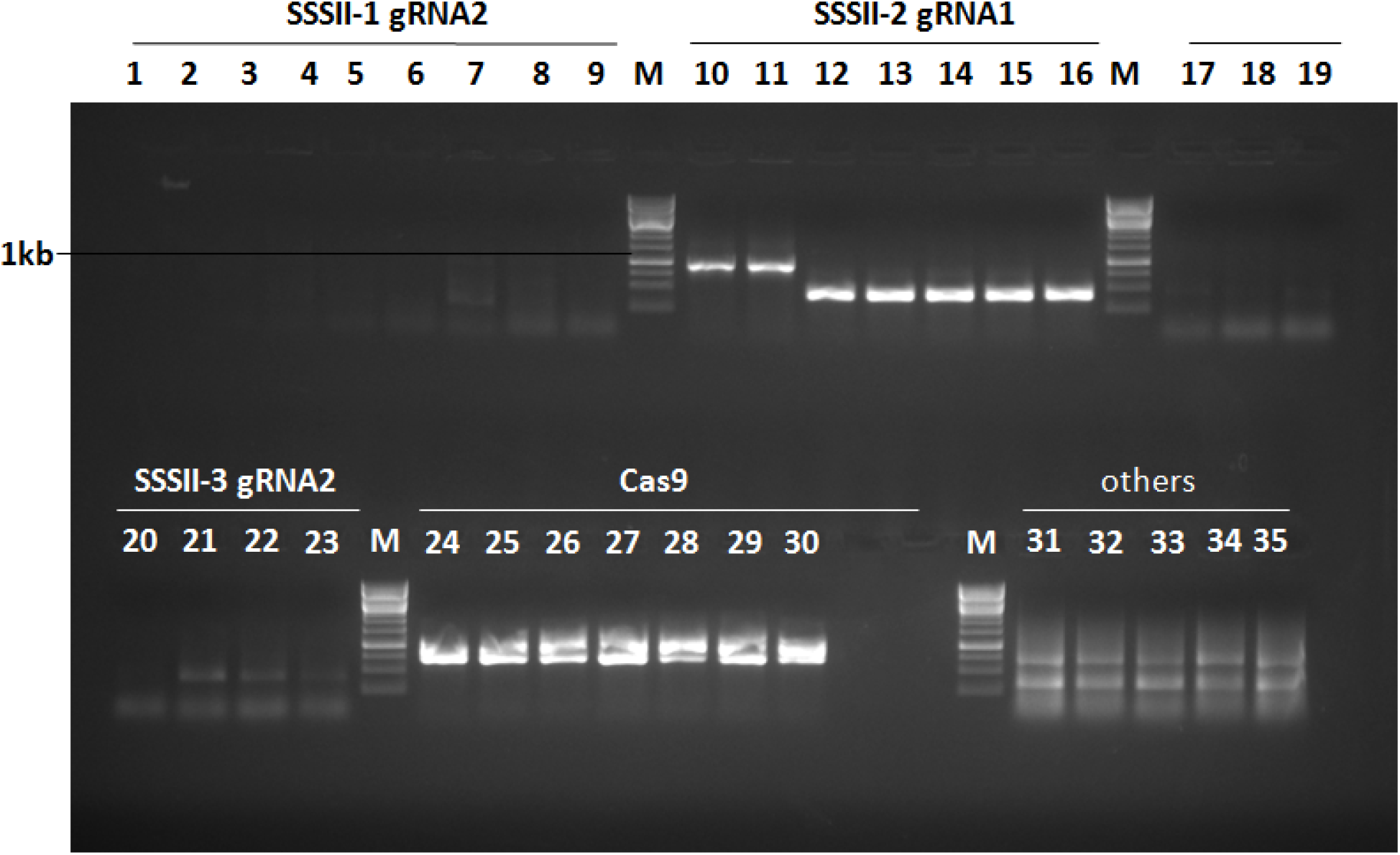
The formation of advance construction of all three isoforms of soluble starch synthases: SSSII-1, 2, and 3 in single destination vector. The Band M is 1 kb DNA ladder. The confirmation with PCR of bands 1 to 9 for SSSII-1 target 2 primers, bands 10 to 16 for SSSII-2 target 1 primers, bands 17 to 23 for SSSII-3 target 2 primers and bands 24 to 30 for Cas9 primers.

**Figure 15:**
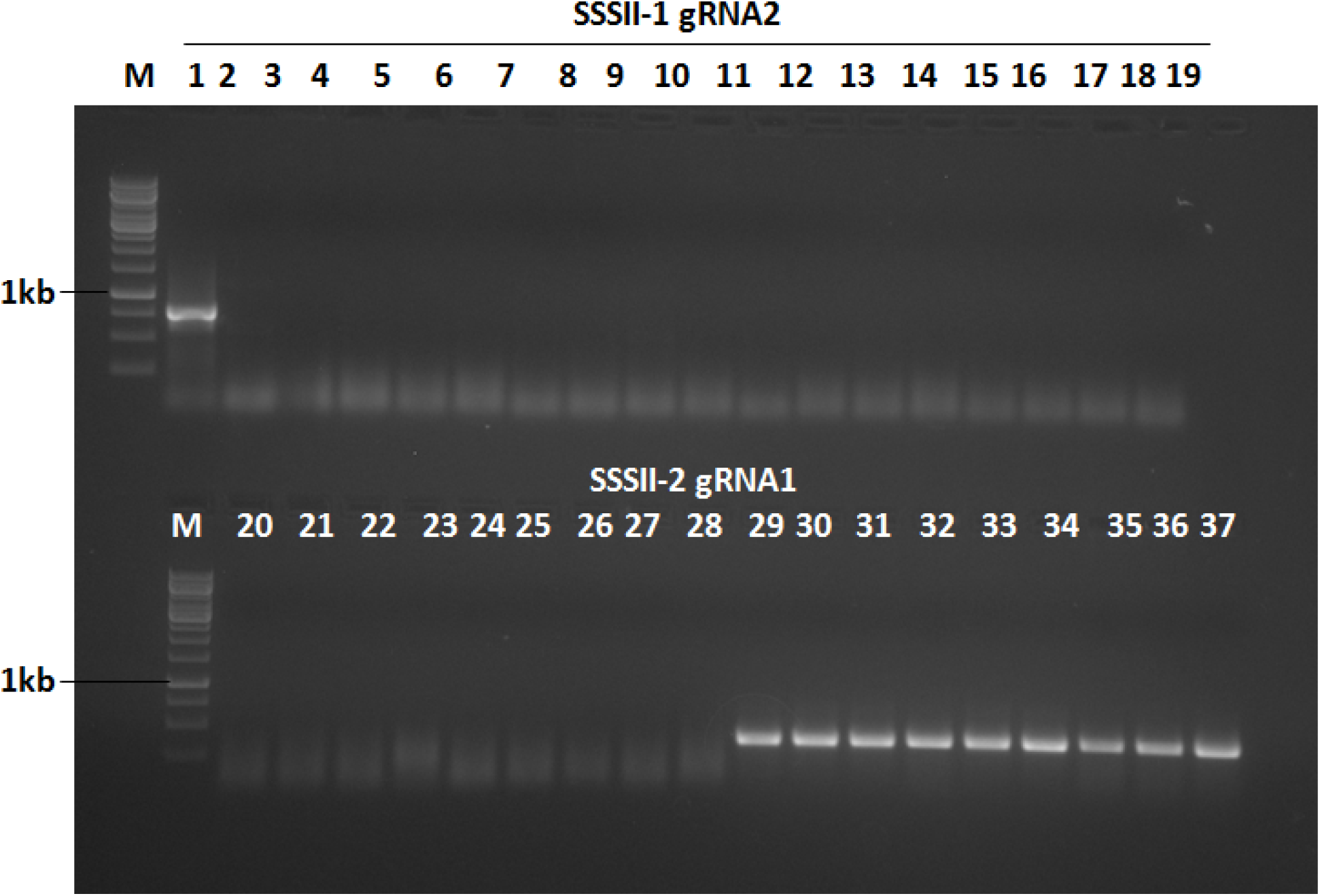
The formation of advance construction of all three isoforms of soluble starch synthases: SSSII-1, 2, and 3 in single destination vector. The Band M is 1 kb DNA ladder. The confirmation with PCR of bands 1 to 19 for SSSII-1 target 2 primers and bands 20 to 37 for SSSII-2 target 1 primers.

**Figure 16:**
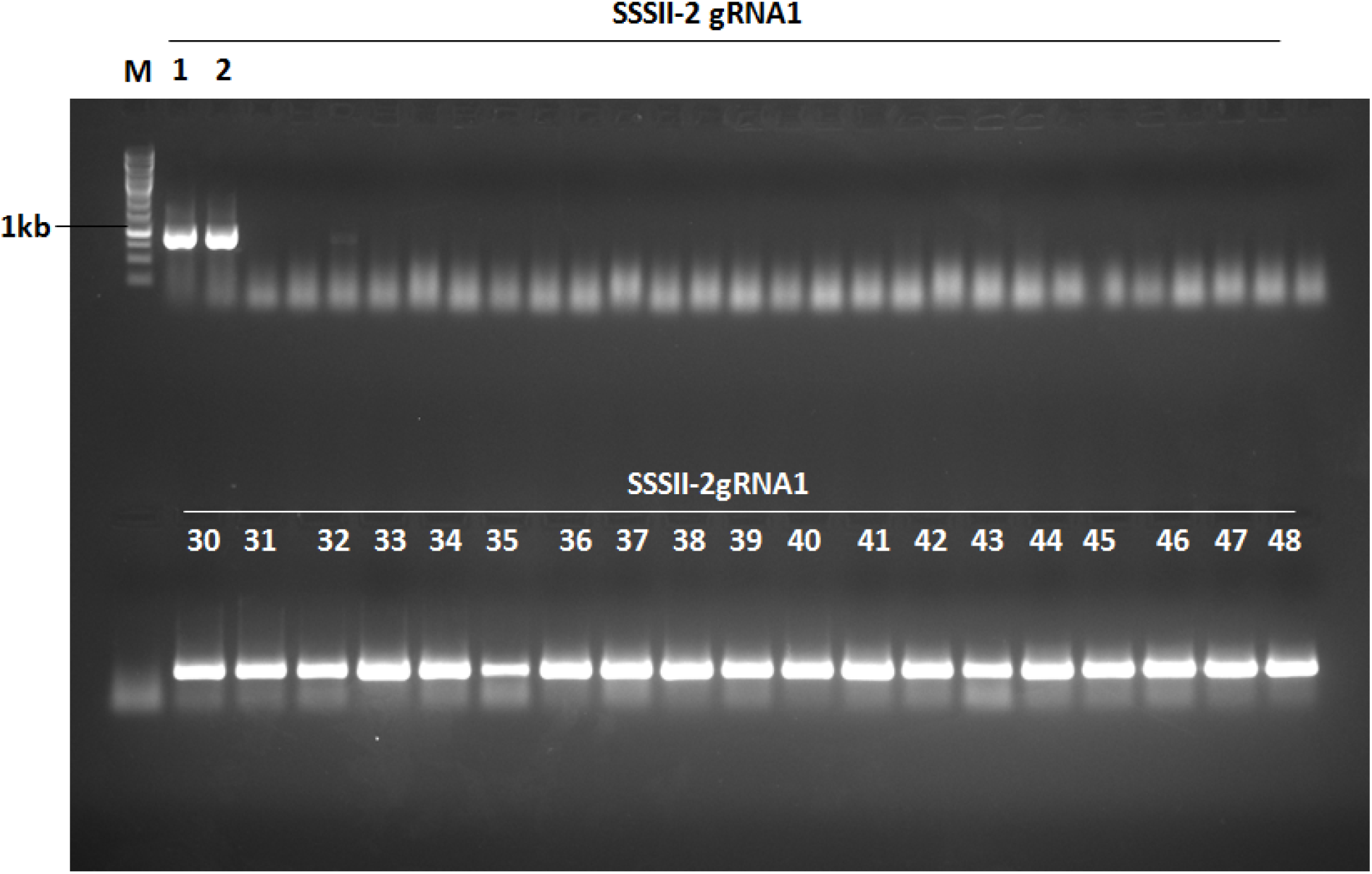
The formation of advance construction of all three isoforms of soluble starch synthases: SSSII-1, 2, and 3 in single destination vector. The Band M is 1 kb DNA ladder. The confirmation with PCR of bands 1 to 48 for SSSII-2 target 2 primers.

**Figure 17:**
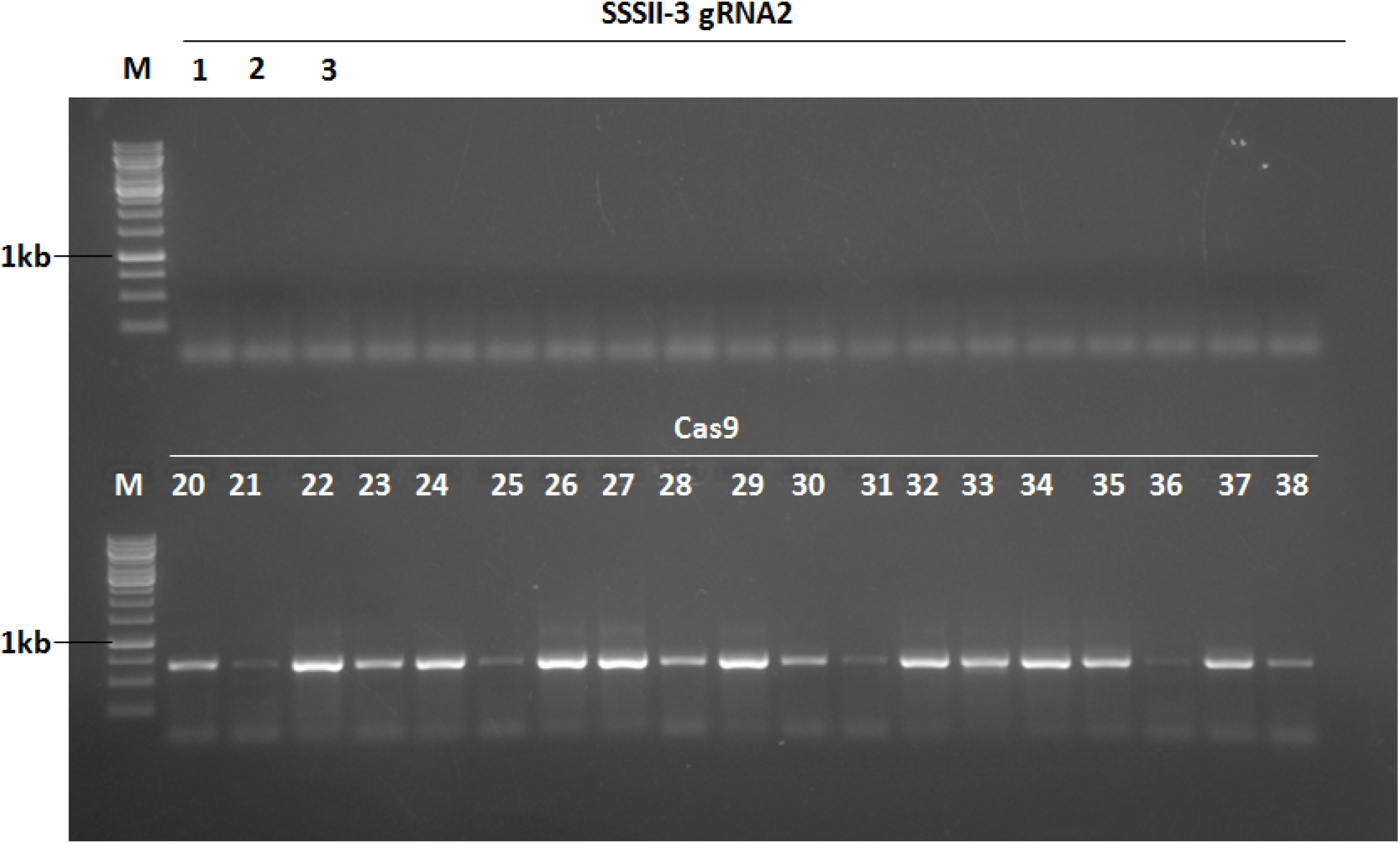
The formation of advance construction of all three isoforms of soluble starch synthases: SSSII-1, 2, and 3 in single destination vector. The Band M is 1 kb DNA ladder. The confirmation with PCR of bands 1 to 19 for SSSII-3 target 2 primers and bands 20 to 38 for SSSII-2 target 1 primers.

### 3. Confirmation of ultimate Binary Expression Vector with *hpt,* Cas9 gene, and the gRNA1, gRNA2 of advance Construct of SSSIIs genes

The LR Multi-Round Gateway cloning was productively utilized to pyramid the gRNAs into the Binary Expression Vector pMDC99. A construct for *SSSII-1, SSSII-2, and SSS II-3* was effectively transformed into *Agrobacterium tumefaciens* (EHA105) through electroporation for consequent transformations of rice. *Agrobacterium* cells harboring the Destination Vector (generous along the gRNAs) were subjected to choice on YEM plates containing the antibiotics rifampicin (10mg/ml), kanamycin (50 mg/ml), and streptomycin (25mg/ml). Then the antibiotic-resistant colonies growing among the plate were screened by colony PCR persecution specific primer sets for Cas9 promoter, U3 and U6 promoter, and gRNAs. Colony PCR of expression vector for *SSSII-1, 2 and 3*(DV; SSSII-1, 2 and 3-gRNA1 and gRNA2) harboring *hpt* factor, Cas9 gene, *SSSII-2*-gRNA1, *SSSII-1*-gRNA2 and *SSSII-3*-gRNA2 gave expected size bands of hpt (~750 bp), Cas9 (~750 bp), gRNA1 (~670 bp), gRNA2 (~370 bp) are shown in figures 4 to 17. Authentication through PCR analysis was found to be necessary to induce positive regarding the integrity of the construct.

### 4. Analysis results of confirmation of Construct of advance construct of SSSIIs genes

In figure 5, the LR third round of Gate cloning is cloning after that conformation of isolation of Plasmid of recombinant Binary vector with gene of primers of *SSSII-1* gRNA2, *SSSII-2* gRNA1, Cas9 and *hpt* in turn of *SSSII-2* gRNA1 bands are not shown in this gel. In figure 6, the LR third round of Gate cloning after that repeats conformation of isolation of Plasmid of recombinant Binary vector with gene of primers of *SSSII-1* gRNA2, *SSSII-2* gRNA1, Cas9 and *hpt* in turn of *SSSII-2* gRNA1 bands are not shown in this gel. In figure 5 and 6, the PCR conformation gel bands of *SSSII-2* gRNA1 in figure 5 and 6 is negative, and further separately confirmation of isolated plasmid of *SSSII-2* gRNA1 is shown positive bands. In figure 14, the LR final round of Multi Gateway cloning for remove ccdB from recombinant binary vectors after that repeats conformation of isolation of Plasmid of recombinant Binary vector with gene of primers of *SSSII-1* gRNA2, *SSSII-1* gRNA2, and *SSSII-3* gRNA2 and Cas9. There are not shown bands of SSSII-1 and SSSII-3. In figure 15, the LR final round of Multi Gateway cloning for remove ccdB from recombinant binary vectors after that repeats conformation of isolation of Plasmid of recombinant Binary vector with gene of primers of *SSSII-1* gRNA2 and *SSSII-1* gRNA2. In figure 16, the LR final round of Multi Gateway cloning for remove ccdB from recombinant binary vectors after that repeats conformation of isolation of Plasmid of recombinant Binary vector with gene of primers of *SSSII-1* gRNA2. In figure 17, the LR final round of Multi Gateway cloning for remove ccdB from recombinant binary vectors after that repeats conformation of isolation of Plasmid of recombinant Binary vector with gene of primers of *SSSII-3* gRNA2 and Cas9.

## DISCUSSION

The CRISPR/Cas9 application is used for gene editing. Gene editing in the intronic region will have no impact. Targeting exon regions is essential for effective approach to delete functional segment of gene in CRISPR/Cas9-mediated genome editing. The gRNA can be changed by digesting the scaffold vector with an appropriate restriction enzyme to clone sgRNA. cDNA sequences of three OsSSSII-1, OsSSSII-2, and OsSSSII-3 genes were used in the CRISPRdirect program predict potential gRNAs. The CRISPRdirect identified few target sites which were promptly adjoining PAM sequences while checking the sense and antisense strands of both the cDNAs. The target sequence resembled 5’-N (20)- NGG-3’ or 5’CCN-N (20)- 3’. The target sequence which is 20 ntds, was available only near the PAM sequence which is 5’NGG3’. Design of primers of CRISPR/Cas9 gRNA using CRISPRdirect is simple and effective [6]. CRISPR-Cas9-targeted mutagenesis has recently been used for creating various mutations in rice [34, 35]. CRISPR/Cas9-mediated genome-edited technology has a major edge over other mutagenesis techniques due to its high precision [36, 12]. Advance Constructing the Gateway technology compatible destination vector. In arrange different genes in advance construct of Gateway cloning compatible binary destination vector. The LR last round recombination consisted of the recombination reaction between SSSII-3 EV2 recombinant vector (having the gRNA2 expression cassette: U6 promoter: gRNA2: scaffold) and the pMDC99:Cas9:gRNA2:gRNA1 vector. The LR final round recombination consisted of the recombination reaction between EV1 vector (having the gRNA1 expression cassette) and the pMDC99:Cas9:gRNA2:gRNA1:gRNA2 vector to remove ccdB gene from advance vector of pMDC99:Cas9: SSSII-1 gRNA2: SSSII-2 gRNA1: SSSII-3 gRNA2. This LR reaction was terminated by adding one μl enzyme K. associate degree aliquot of 2μl of the primary round LR reaction product was remodeled into E. coli top10/ E. coli DB3.1 (EV2/gRNA2 is related to E. coli DB3.1) cells exploitation heat shock and designated on lb agar plates supplemented with 50 μg/ml kanamycin. The resistant colonies obtained were screened for the presence of a gRNAs expression cassette by colony PCR exploitation sequence-specific primers. The LR final round cloning was transferred into *Agrobacterium* (EHA105) via electroporation and chosen on YEM agar plate supplemented with 50 μg/ml kanamycin, 50 μg/ml chloramphenicol and 50 μg/ml rifampicin. The ultimate expression vector construct (pMDC99:Cas9: OsSSSII-1-gRNA1:gRNA2, pMDC99: Cas9: OsSSSII-2-gRNA1: gRNA2 and pMDC99: Cas9: OsSSSII-3-gRNA1: gRNA2) was checked for the presence of *hpt* gene, Cas9 gene, gRNA1, and gRNA2 discrimination specific sets of screening primers. Delineated illustration of ultimate expression vector having Cas9, gRNA expression cassette. The sequence of screening primers of hygromycin, Cas9 genes, gRNAs of all three *OsSSSII-1, OsSSSII-2 and OsSSSII-3* Expression. The LR third round of Gate cloning is cloning after that conformation of isolation of Plasmid of recombinant Binary vector with gene of primers of SSSII-1 gRNA2, SSSII-2 gRNA1, Cas9 and *hpt,* but SSSII-2 gRNA1 bands are not shown in this gel (figure 5). This is repeats conformation of isolation of Plasmid. In figure 5 and 6, the PCR conformation gel bands of SSSII-2 gRNA1 in figure 5 and 6 is negative, and further separately confirmation of isolated plasmid of SSSII-2 gRNA1 is shown positive bands. The LR final round of Multi Gateway cloning for remove ccdB from recombinant binary vectors after that repeats conformation of isolation of Plasmid of recombinant Binary vector with gene of primers of SSSII-1 gRNA2, SSSII-1 gRNA2, and SSSII-3 gRNA2 and Cas9 (figure 14). There are not shown bands of SSSII-1 and SSSII-3. The LR final round of Multi Gateway cloning for remove ccdB from recombinant binary vectors after that repeats conformation of isolation of Plasmid of recombinant Binary vector with gene of primers of SSSII-1 gRNA2 and SSSII-1 gRNA2 (figure 15). The LR final round of Multi Gateway cloning for remove ccdB from recombinant binary vectors after that repeats conformation with respective gene primers (figure 16 and 17). Unfortunately, we did not get successful for formation of advance construction of all three genes. The rice developed had a low glycemic index and thus offers a good substitute for traditional rice for diabetics and patients suffering from other diseases. Cas9-free SSS mutant rice can help in controlling diabetes, coronary heart disease, rectal cancers, and certain colon cancers, and might offer health benefits for people worldwide [12].

## CONCLUSION

In this Article, CRISPR/Cas9 system is used for editing of targeted sequences of genomes. CRISPR-Cas9 is used for faster and easier modification other than previous genome editing. The formation advances construct CRISPR/Cas9 of soluble starch synthase II of three isoforms gene in the single destination binary vector (pMDC99) with Multi round LR reaction of Gateway cloning technology. In fourth LR reaction, we inserted three gRNAs of gene in binary vector. We tried to remove ccdB gene from construct vector through final LR reaction of gateway technology, unfortunately disturbed construct of three genes of SSSII. We reported results in this article. This technology is used for knock out of targeted sequences.

## AUTHOR CONTRIBUTIONS

Conceptualization, M.R.J. and S.A.; Data curation, S.A.; Formal analysis, M.R.J., F.J. and S.A.; Investigation, S.A.; Methodology, M.R.J., F.J. and S.A.; Project administration, M.M.; Software, M.R.J., S.A. and M.M.; Supervision, S.A.; Validation, M.R.J.,S.A. and M.M.; Visualization, M.M. and F.J.; Writing—original draft, M.R.J.; Writing— review and editing, S.A. and S.A. All authors have read and agreed to the published version of the manuscript.

## ACKNOWLEDGMENTS

We would like to thank Dr. MK Reddy, International Centre for Genetic engineering and Biotechnology, New Delhi, Co-supervisor of M. Rizwan Jameel, for his generous help in this study.

## CONFLICTS OF INTEREST

The authors declare no competing interests.

## Notes

### Competing Interest Statement

The authors have declared no competing interest.

